## Supplementary material for "NF-_κ_B is a Central Regulator of Hypoxia-Induced Gene Expression": suppplementary figures_Appendix

^1^Department of Biochemistry, Cell and System Biology. Institute of Systems, Molecular and Integrative Biology, University of Liverpool. Liverpool L697ZB. United Kingdom.

^2^Signalling Programme, The Babraham Institute, Babraham Research Campus, Cambridge CB22 3AT. United Kingdom.

*Corresponding author

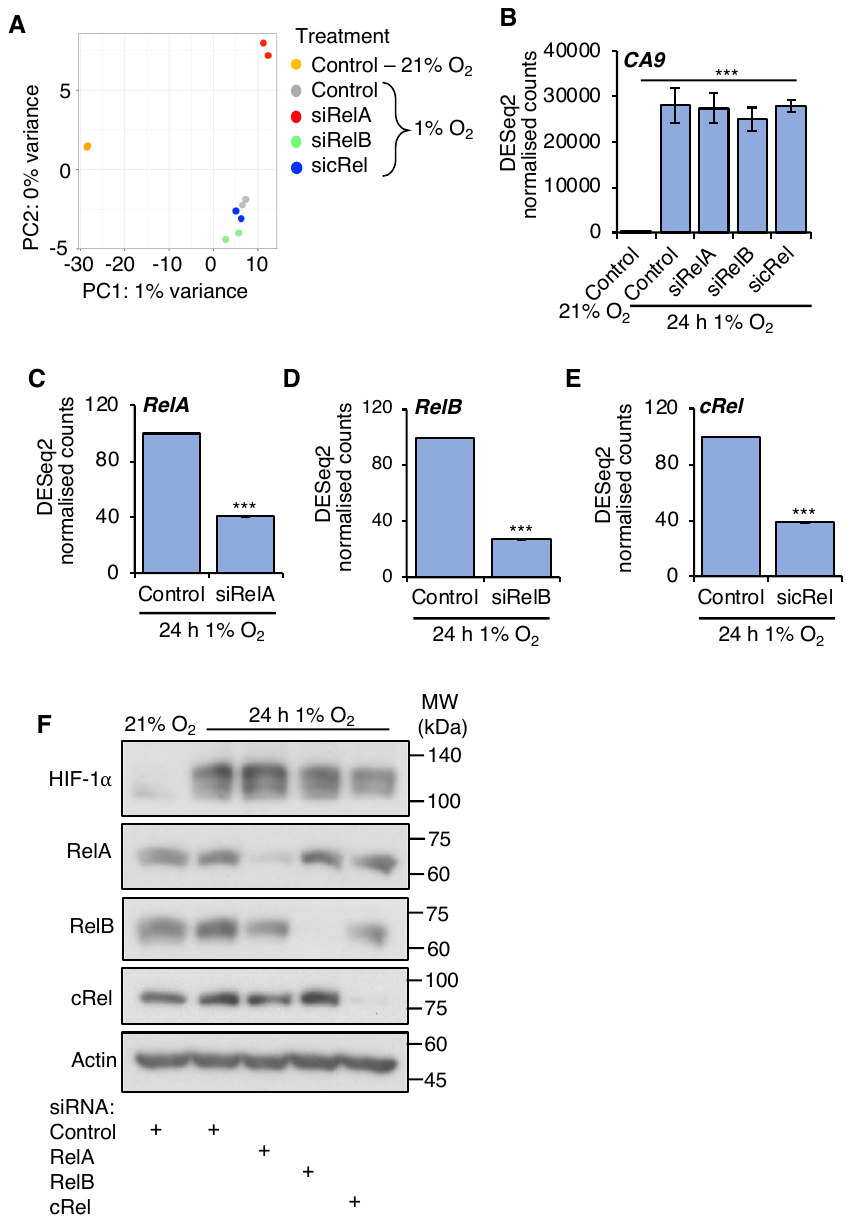
Appendix Figure S1 **Gene expression changes in response to hypoxia and NF-𝜅B subunit depletions, additional data.**

**A.** RNA-seq (n=2) principal component analysis (PCA) sample clustering by treatment. Orange points indicate cells cultured at 21% oxygen (normoxia) and transfected with control siRNA, grey points indicate cells cultured at 24 h 1% oxygen (hypoxia) and transfected with control siRNA, red points indicate cells cultured at hypoxia and transfected with RelA siRNA (siRelA), green points indicate cells cultured at hypoxia and transfected with RelB siRNA (siRelB), blue points indicate cells cultured at hypoxia and transfected with cRel siRNA (sicRel). **B.** DESeq2 normalised counts acquired from RNA-seq analysis showing differentially expressed CA9 gene in hypoxia control, and siRelA, siRelB and sicRel with hypoxia, compared to normoxia control. **C-E.** DESeq2 normalised counts of differentially expressed RelA (**C**), RelB (**D**), and cRel (**E**) genes with siRelA, siRelB and sicRel in hypoxia, compared to hypoxia control. **B-E.** Statistical significance of the gene list overlaps was determined via DESeq2 differential expression analysis, *** FDR < 0.001. **F.** Immunoblot analysis of the indicated proteins in HeLa cells cultured at normoxia or hypoxia, transfected with control siRNA or RelA, RelB or cRel siRNAs. Representative images from 3 independent experiments are shown.

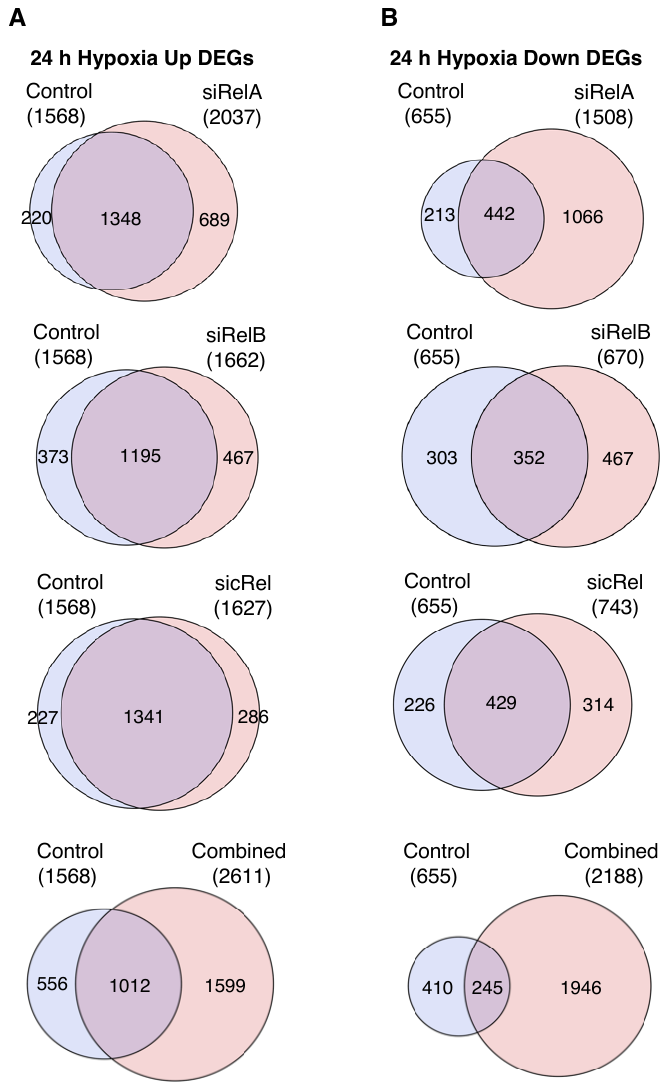

Appendix Figure S2 **Identifying NF-𝜅B dependent hypoxia responsive genes.**

RNA-seq (n=2) in HeLa cells cultured at 21% oxygen (normoxia) or exposed to 24 h 1% oxygen (hypoxia), transfected with control siRNA or RelA, RelB or cRel siRNAs. **A.** Overlap of hypoxia control siRNA upregulated genes (compared to normoxia control siRNA) with hypoxia siRelA, siRelB, sicRel, and the combined list of siRelA, siRelB, and sicRel upregulated genes (compared to normoxia control siRNA). **B.** Overlap of hypoxia control siRNA downregulated genes (compared to normoxia control siRNA) with hypoxia siRelA, siRelB, sicRel, and the combined list of siRelA, siRelB, and sicRel downregulated genes (compared to normoxia control siRNA). **A-B.** Genes which are differentially regulated in hypoxia with control siRNA, but not RelA/RelB/cRel targeting siRNA, are considered RelA/RelB/cRel dependent hypoxia responsive genes.

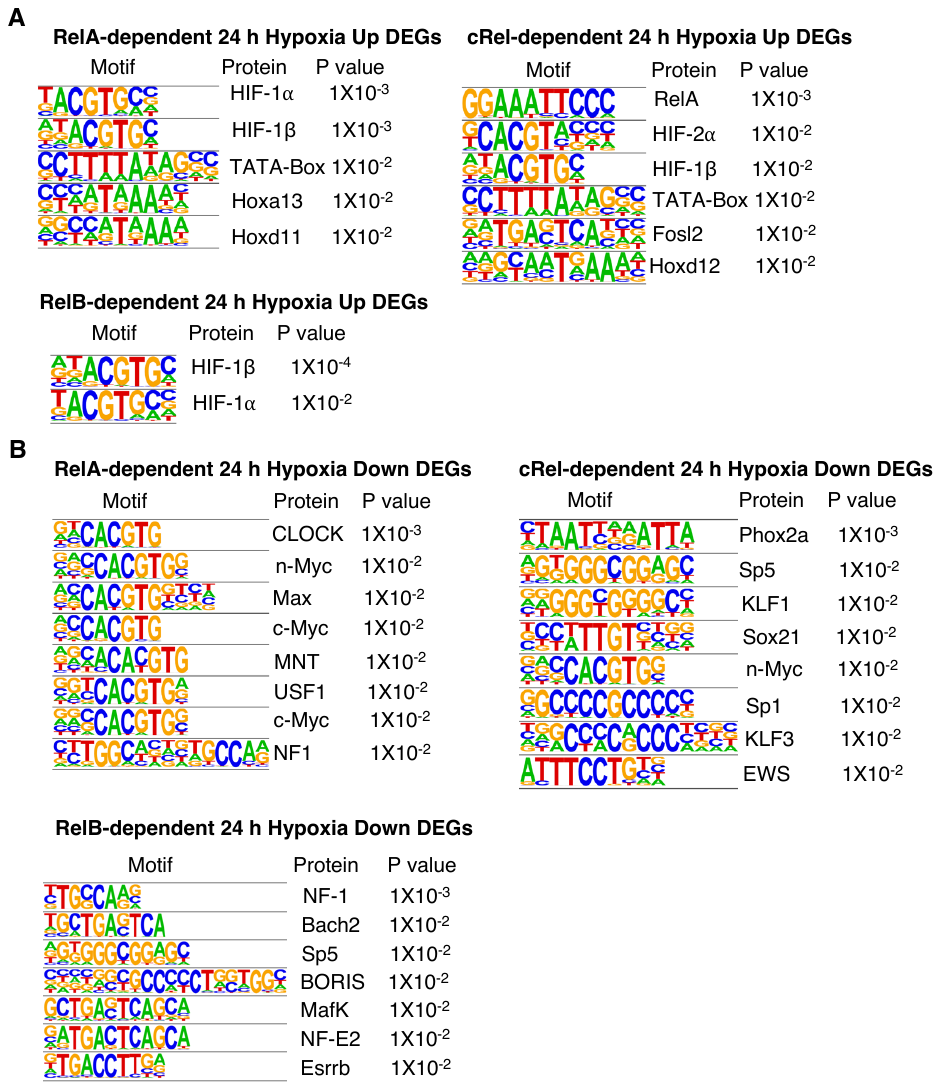

Appendix Figure S3 **Motif enrichment of NF-𝜅B dependent hypoxia inducible DEGs.**

**A-C.** Transcription factor motif enrichment analysis for RelA-, RelB-, and cRel-dependent hypoxia up-regulated genes (**A**), RelA-, RelB-, and cRel-dependent hypoxia down-regulated genes (**B**), and a randomly selected set of 500 hypoxia upregulated genes and 500 hypoxia downregulated genes.

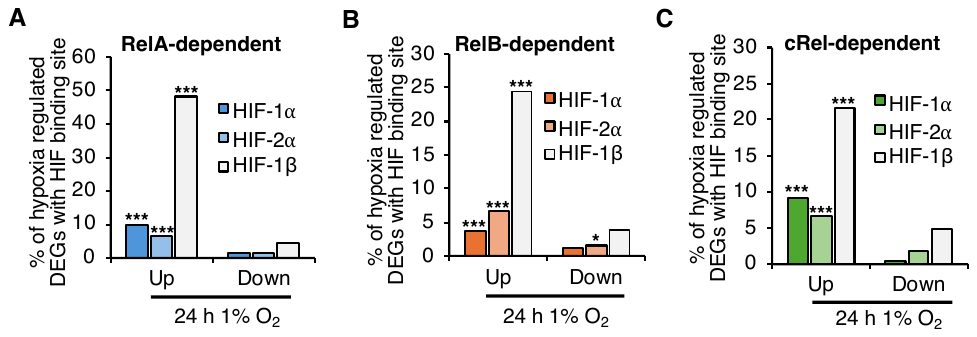
Appendix Figure S4 **Hypoxia inducible factor (HIF) subunit dependency of the gene signature.**

**A-C.** Overlap of NF-𝜅B dependent hypoxia up- and down-regulated DEGs identified by RNA-seq (n=2) with HIF subunit binding sites identified by ChIP-seq analysis (n=2) in HeLa cells exposed to 6 hours (h) hypoxia. Percentage of DEGs containing a HIF binding site (i.e., HIF-1α, HIF-2α or HIF-1β) are displayed; statistical significance was determined via hypergeometric test, *** P < 0.001.

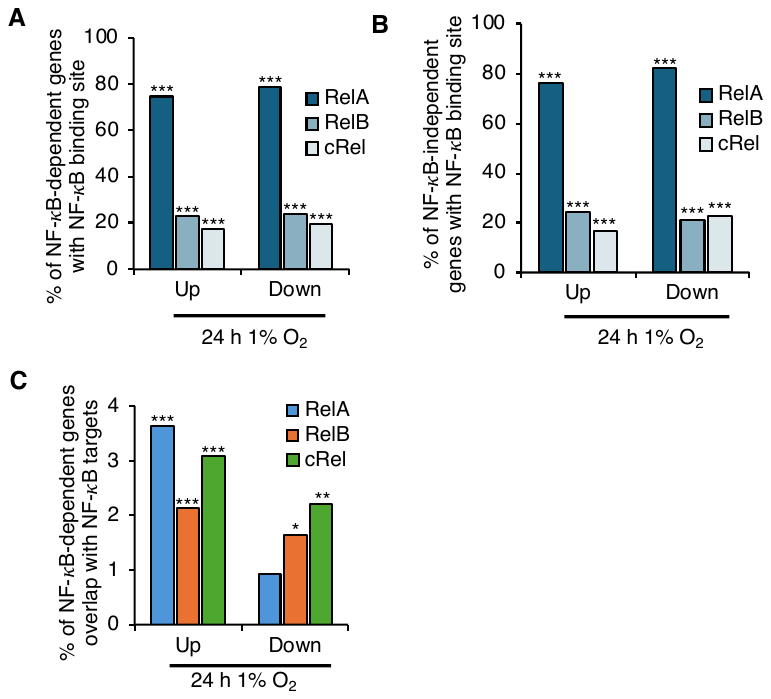
Appendix Figure S5 **Association of NF-𝜅B dependent hypoxia inducible DEGs with NF-𝜅B binding sites and target genes.**

**A-B.** Overlap of NF-𝜅B dependent (A) and independent (B) hypoxia up- and down-regulated DEGs identified by RNA-seq (n=2) with NF-𝜅B subunit binding sites acquired from the ChIP-seq Atlas database. Percentage of DEGs containing an NF-𝜅B subunit binding site are displayed. **C.** Percentage values for the overlap of NF-𝜅B dependent hypoxia inducible DEGs identified in HeLa cell RNA-seq dataset with Gilmore laboratory’s NF-𝜅B target genes; statistical significance was determined via hypergeometric test, * P < 0.05 ** P < 0.01, *** P < 0.001**.**

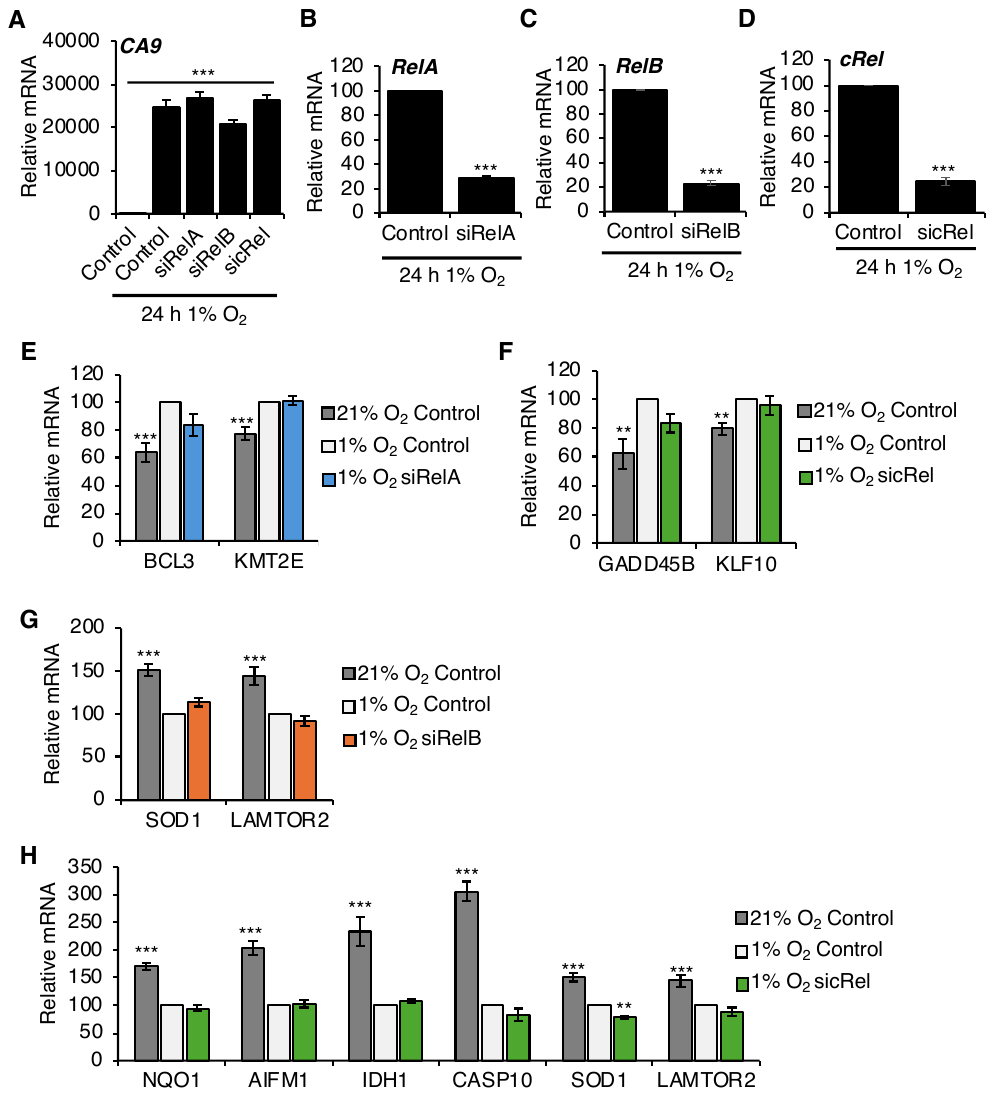
Appendix Figure S6 **Validation of NF-𝜅B dependent hypoxia inducible gene expression changes in HeLa cells, additional data.**

**A-D.** qPCR analysis showing *CA9* gene (**A**) expression in hypoxia control (24 h 1% O_2_), and siRelA, siRelB and sicRel with hypoxia, compared to normoxia control (21% O_2_); *RelA* (**B**), *RelB* (**C**), and *cRel* (**D**) gene expressions with depletion of each NF-𝜅B subunits in hypoxia, compared to hypoxia control. **E-F.** Hypoxia upregulated DEG changes with RelA- (**E**) or cRel- (**F**) depletion in hypoxia. **G-H.** Hypoxia downregulated DEG changes with RelB- (**G**) or cRel- (**H**) depletion in hypoxia. Graphs show mean (n = 4) ± SEM, * P < 0.05, ** P < 0.01, *** P < 0.001. Statistical significance was determined via one-way ANOVA with post-hoc Dunnett’s test for comparing more than two conditions. Student’s t-test was applied while comparing two conditions.

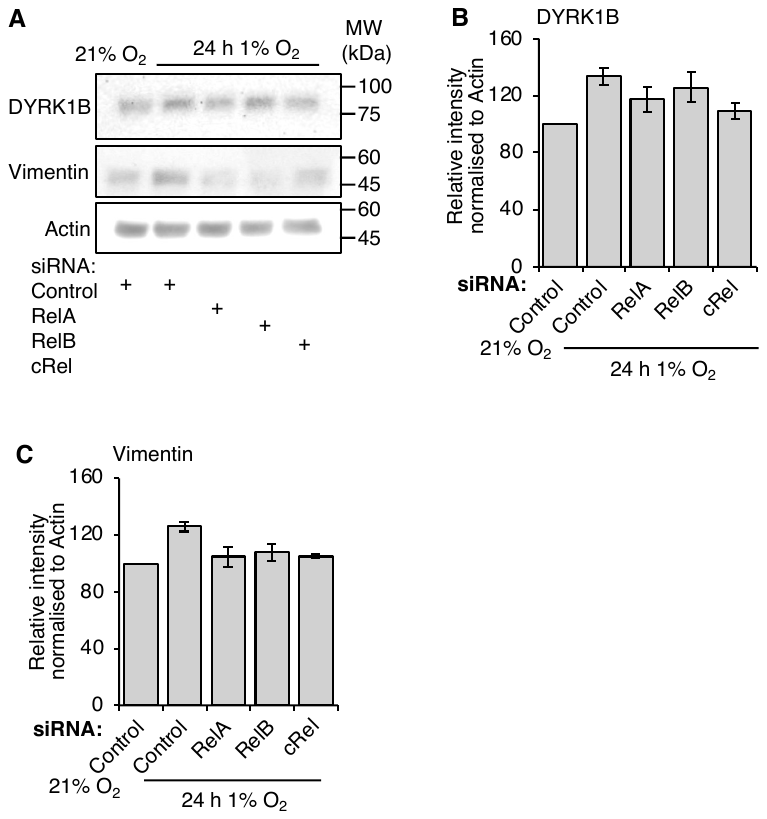

Appendix Figure S7 **Validation of NF-𝜅B dependent hypoxia inducible gene expression changes in HeLa cells at protein level.**

**A.** Immunoblot analysis of the indicated proteins in HeLa cells exposed or not to 1% oxygen (hypoxia) for 24 h, with siRNA transfection of control, RelA, RelB or cRel. Representative images from 3 independent experiments are shown. **B-C.** Signal intensity normalised to Actin (mean n=3, ± SEM) from immunoblot analysis.

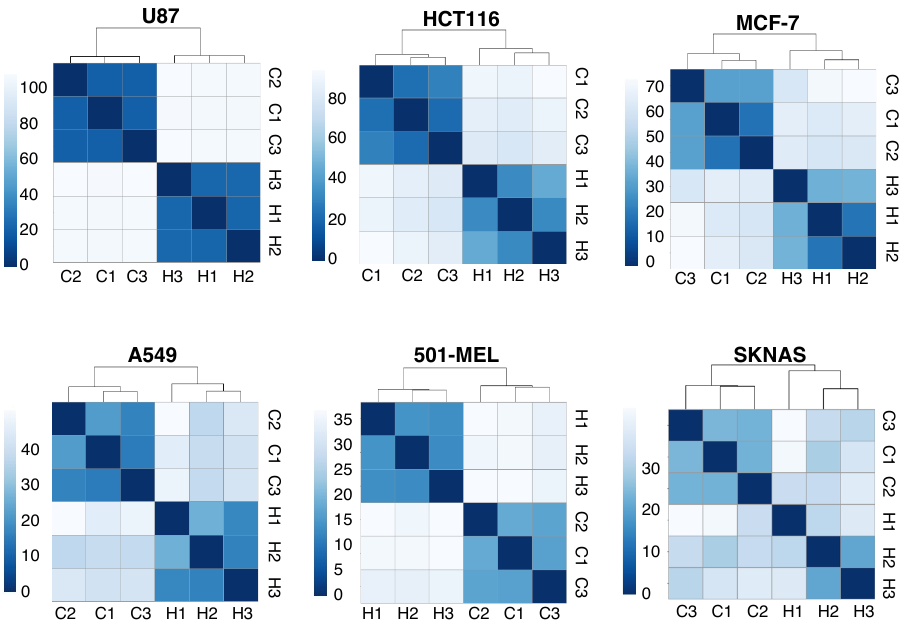
Appendix Figure S8 **Identification of NF-𝜅B dependent hypoxia inducible gene signature in different cell types, additional data.**

Sample to sample distance heatmaps of normalised read counts with hierarchical clustering of RNA-sequencing samples in each of the indicated cell lines. Control (C) represents normoxic (21% O_2_) samples, Hypoxia (H) represents hypoxic (24 hours, 1% O_2_) samples.

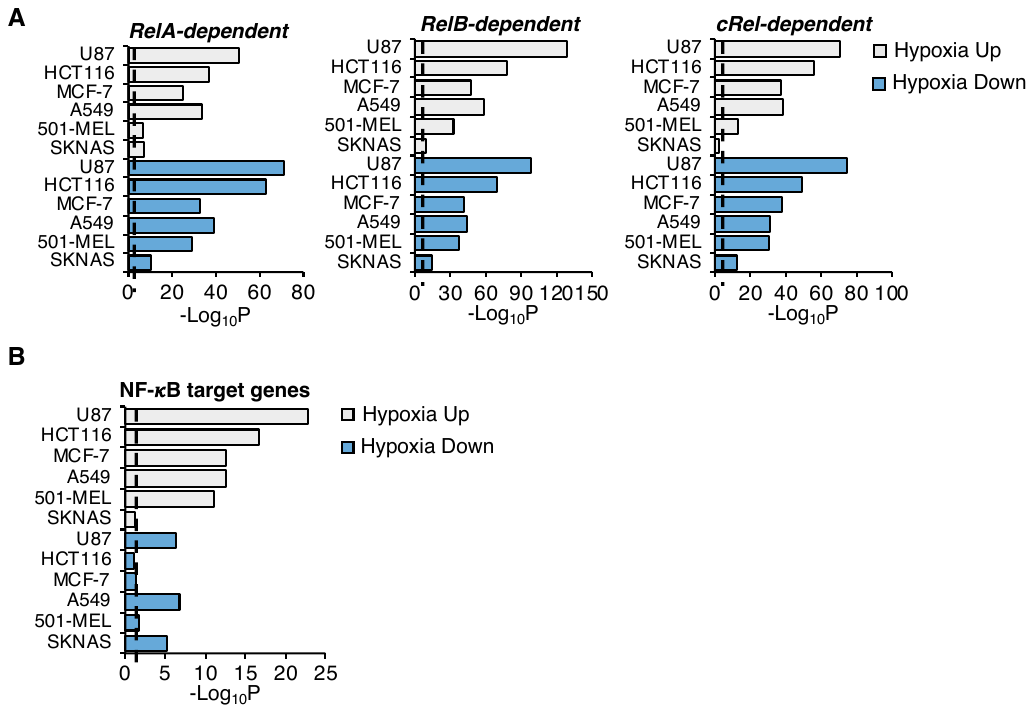

Appendix Figure S9 **Identification of NF-𝜅B dependent hypoxia inducible gene signature in different cell types, additional data.**

**-**Log_10_P values for the overlap of hypoxia inducible DEGs identified in different cell lines’ RNA-seq datasets with NF-𝜅B dependent hypoxia inducible DEGs identified in HeLa RNA-seq experiment following siRNA depletion of individual NF-𝜅B subunits exposed or not to 24 h hypoxia (**A**). **-**Log_10_P values for the overlap of hypoxia inducible DEGs identified in different cell lines’ RNA-seq datasets with Gilmore laboratory’s NF-𝜅B target genes (**B**). Dashed line shows statistical significance threshold of P value 0.05 (-log10 P 1.3). Statistical significance of the gene list overlaps was determined via hypergeometric test, * P < 0.05, ** P < 0.01, *** P < 0.001.

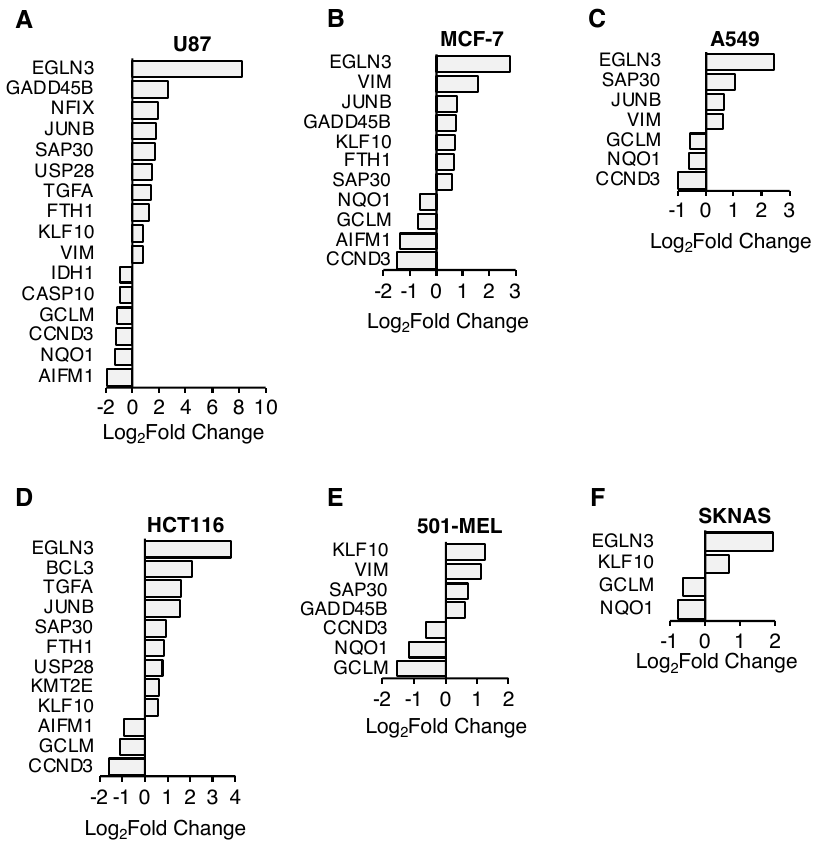

Appendix Figure S10 **Identification of NF-𝜅B dependent hypoxia induced DEGs in various cellular backgrounds.**

**A-F.** Log_2_Fold change of selected NF-𝜅B dependent hypoxia up- and down-regulated DEGs, showing altered transcript levels in hypoxia compared to normoxia identified by RNA-seq (n=3) in different cell lines.

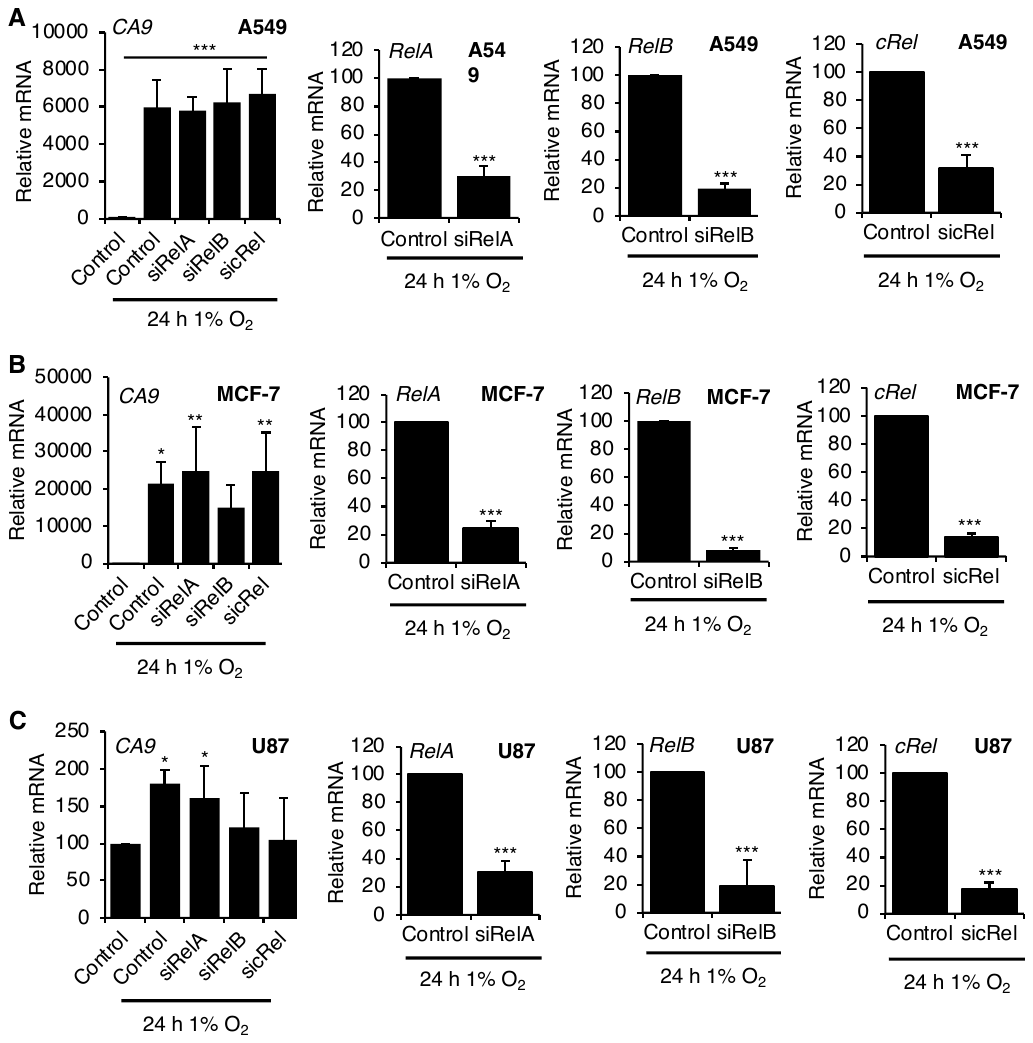
Appendix Figure S11 **Validation of NF-𝜅B dependent hypoxia inducible gene expression changes in HeLa cells, additional data.**

**A-C.** qPCR analysis showing *CA9* gene expression in hypoxia control (24 h 1% O_2_), and siRelA, siRelB and sicRel with hypoxia, compared to normoxia control (21% O_2_); and *RelA*, *RelB*, and *cRel* gene expressions with depletion of each NF-𝜅B subunits in hypoxia, compared to hypoxia control in different cell lines. Graphs show mean (n = 3) ± SEM, * P < 0.05, ** P < 0.01, *** P < 0.001. Statistical significance was determined via one-way ANOVA with post-hoc Dunnett’s test for comparing more than two conditions. Student’s t-test was applied while comparing two conditions.

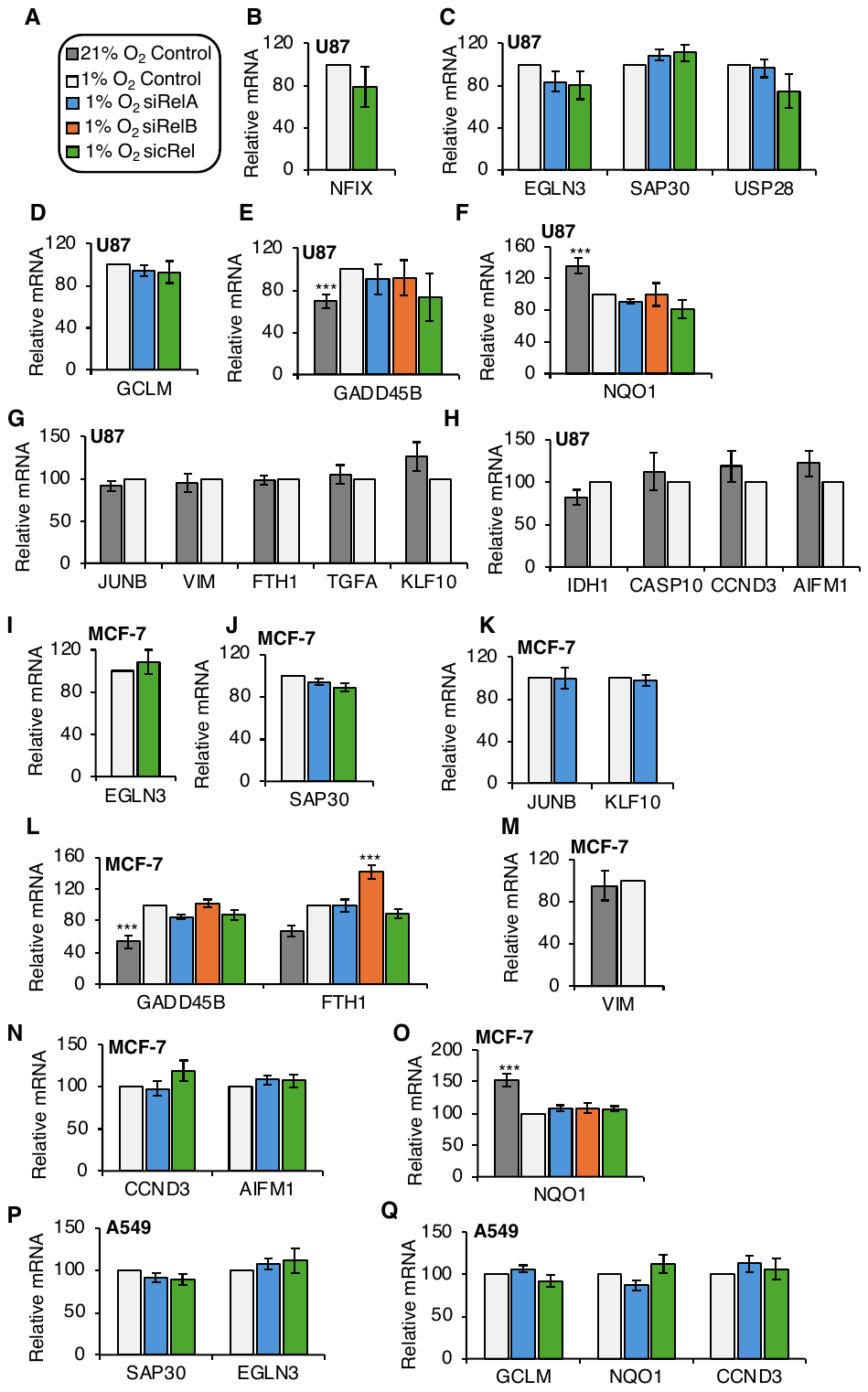

Appendix Figure S12 **Analysis of hypoxia inducible NF-𝜅B-dependent genes across various cell backgrounds, additional data. A.** Key for qPCR analysis graphs. **B-F.** mRNA levels unchanged genes with NF-𝜅B depletions in hypoxia in U87 cells. **G-H.** mRNA levels unchanged genes in hypoxia in U87 cells. **I-O.** mRNA levels unchanged genes with NF-𝜅B depletions in hypoxia in MCF-7 cells. **P-Q.** mRNA levels unchanged genes with NF-𝜅B depletions in hypoxia in A549 cells. Relative mRNA expression levels of the indicated genes were analysed using Actin as a normalising gene for U87 and MCF-7 cell lines (**B-O**), and 18S for A549 cells (**P-Q**). Graphs show mean (n=3) ± SEM, * P < 0.05, ** P < 0.01, *** P < 0.001. Statistical significance was determined via one-way ANOVA with post-hoc Dunnett’s test.

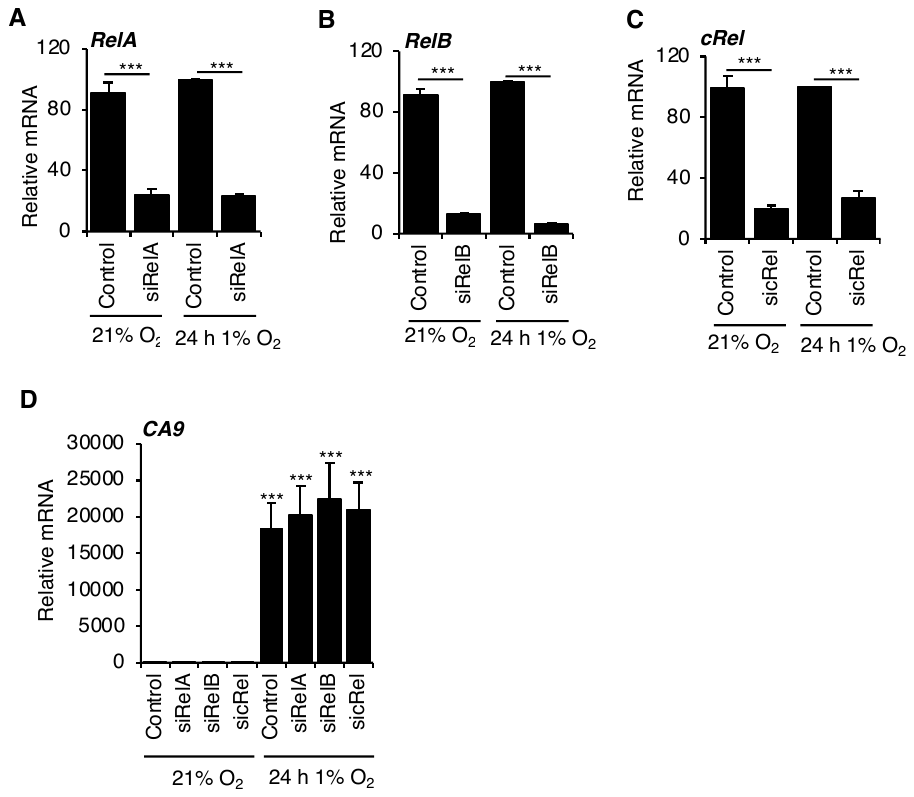
Appendix Figure S13 **Controls for NF-𝜅B dependent gene expression changes in HeLa cells in normoxia and hypoxia. A-D.** qPCR analysis showing, *RelA*, *RelB*, *cRel* and *CA9* gene expression in HeLa cells cultured at 21% oxygen (normoxia) or 24 h 1% oxygen (hypoxia), transfected with control siRNA or RelA, RelB or cRel siRNAs. Relative mRNA expression levels were analysed using Actin as a normalising gene. Graphs show mean (n=3) ± SEM, * P < 0.05, ** P < 0.01, *** P < 0.001. **A-C.** Statistical significance was determined via one-way ANOVA with post-hoc Tukey's HSD test. **D.** Statistical significance was determined via one-way ANOVA with post-hoc Dunnett’s test (compared to normoxia control siRNA treated cells).

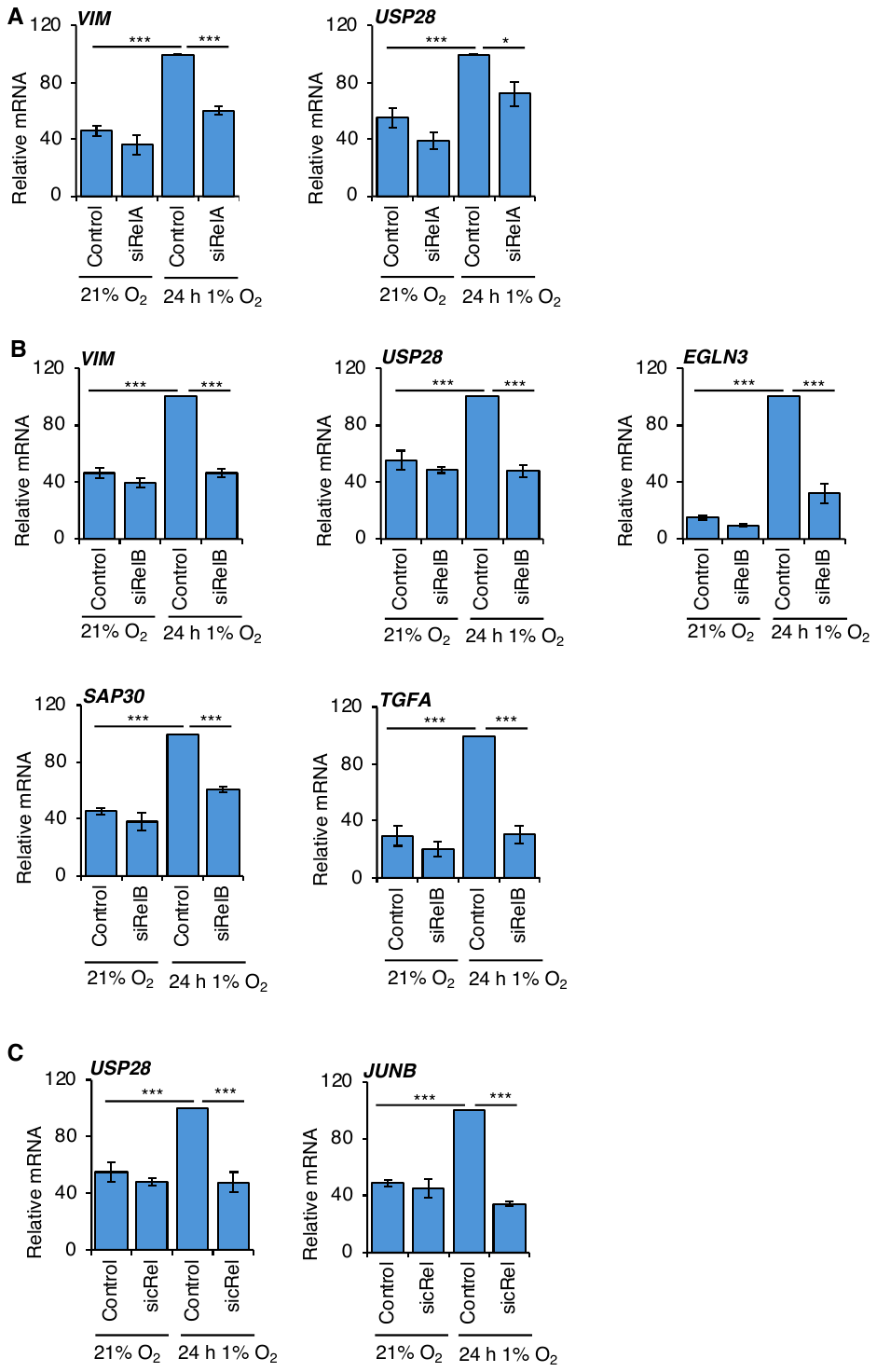

Appendix Figure S14 **NF-𝜅B dependent gene expression changes in HeLa cells in normoxia and hypoxia. A-C.** qPCR analysis in HeLa cells cultured at 21% oxygen (normoxia) or 24 h 1% oxygen (hypoxia), transfected with control siRNA or RelA, RelB or cRel siRNAs. RelA- (**A**), RelB- (**B**), and cRel-dependent (**C**) hypoxia DEGs. Relative mRNA expression levels of the indicated genes were analysed using a as a normalising gene. Graphs show mean (n=3) ± SEM, * P < 0.05, ** P < 0.01, *** P < 0.001. Statistical significance was determined via one-way ANOVA with post-hoc post-hoc Tukey's HSD test.

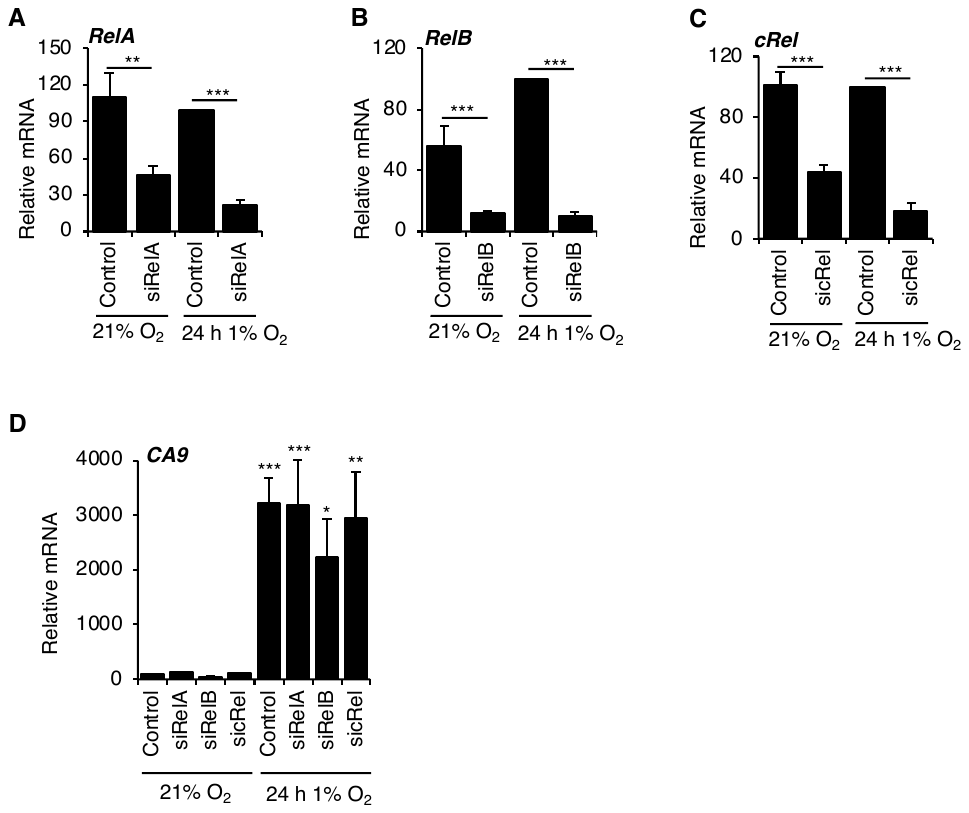

Appendix Figure S15 **Controls for NF-𝜅B dependent gene expression changes in A549 cells in normoxia and hypoxia. A-C.** qPCR analysis showing, *RelA*, *RelB*, *cRel* and *CA9* gene expression in A549 cells cultured at 21% oxygen (normoxia) or 24 h 1% oxygen (hypoxia), transfected with control siRNA or RelA, RelB or cRel siRNAs. Relative mRNA expression levels were analysed using Actin as a normalising gene. Graphs show mean (n=3) ± SEM, * P < 0.05, ** P < 0.01, *** P < 0.001. **A-C.** Statistical significance was determined via one-way ANOVA with post-hoc Tukey's HSD test. **D.** Statistical significance was determined via one-way ANOVA with post-hoc Dunnett’s test (compared to normoxia control siRNA treated cells).

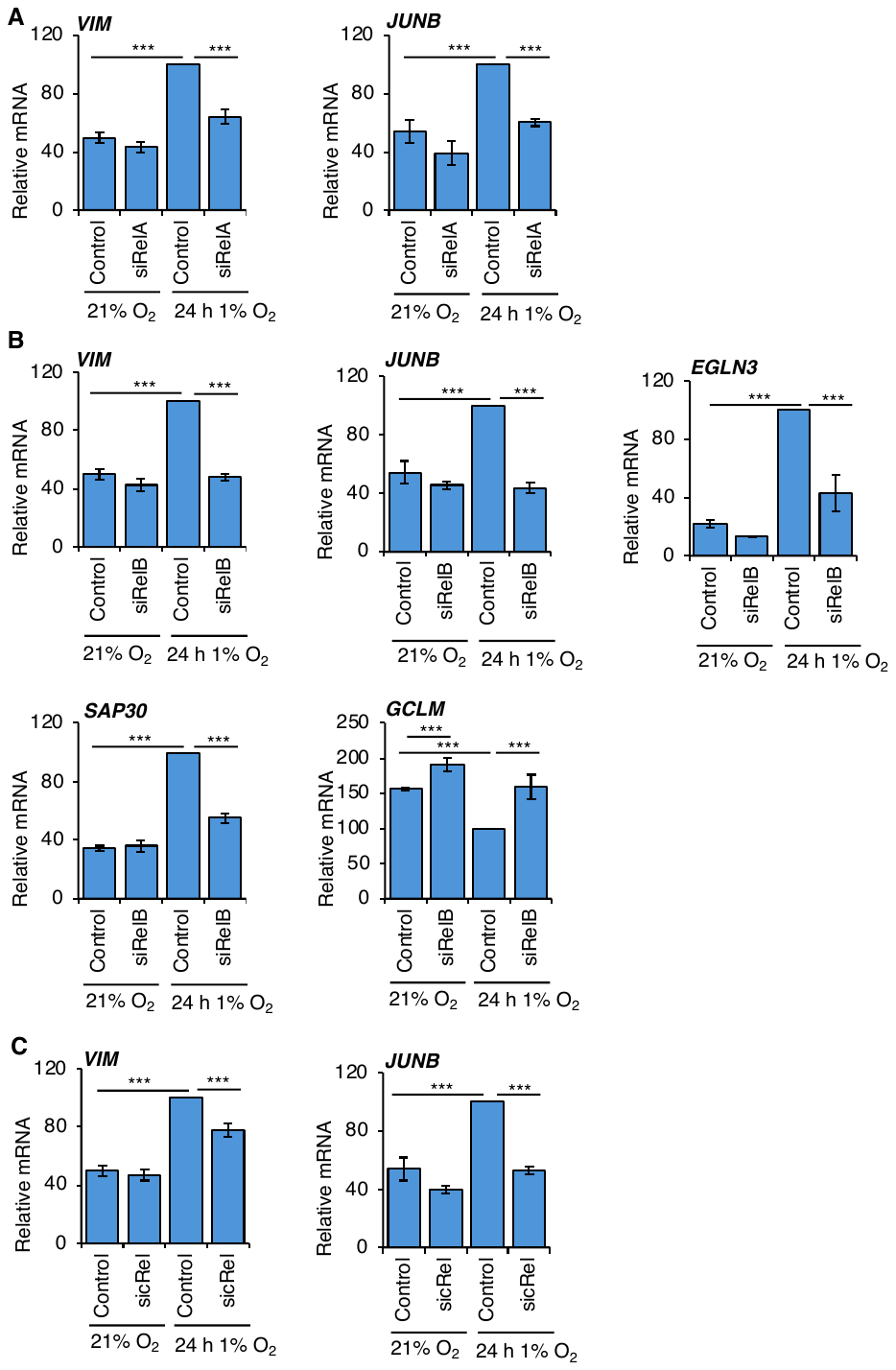
Appendix Figure S16 **NF-𝜅B dependent gene expression changes in A549 cells in normoxia and hypoxia. A-C.** qPCR analysis in A549 cells cultured at 21% oxygen (normoxia) or 24 h 1% oxygen (hypoxia), transfected with control siRNA or RelA, RelB or cRel siRNAs. RelA- (**A**), RelB- (**B**), and cRel-dependent (**C**) hypoxia DEGs. Relative mRNA expression levels of the indicated genes were analysed using a as a normalising gene. Graphs show mean (n=3) ± SEM, * P < 0.05, ** P < 0.01, *** P < 0.001. Statistical significance was determined via one-way ANOVA with post-hoc post-hoc Tukey's HSD test.

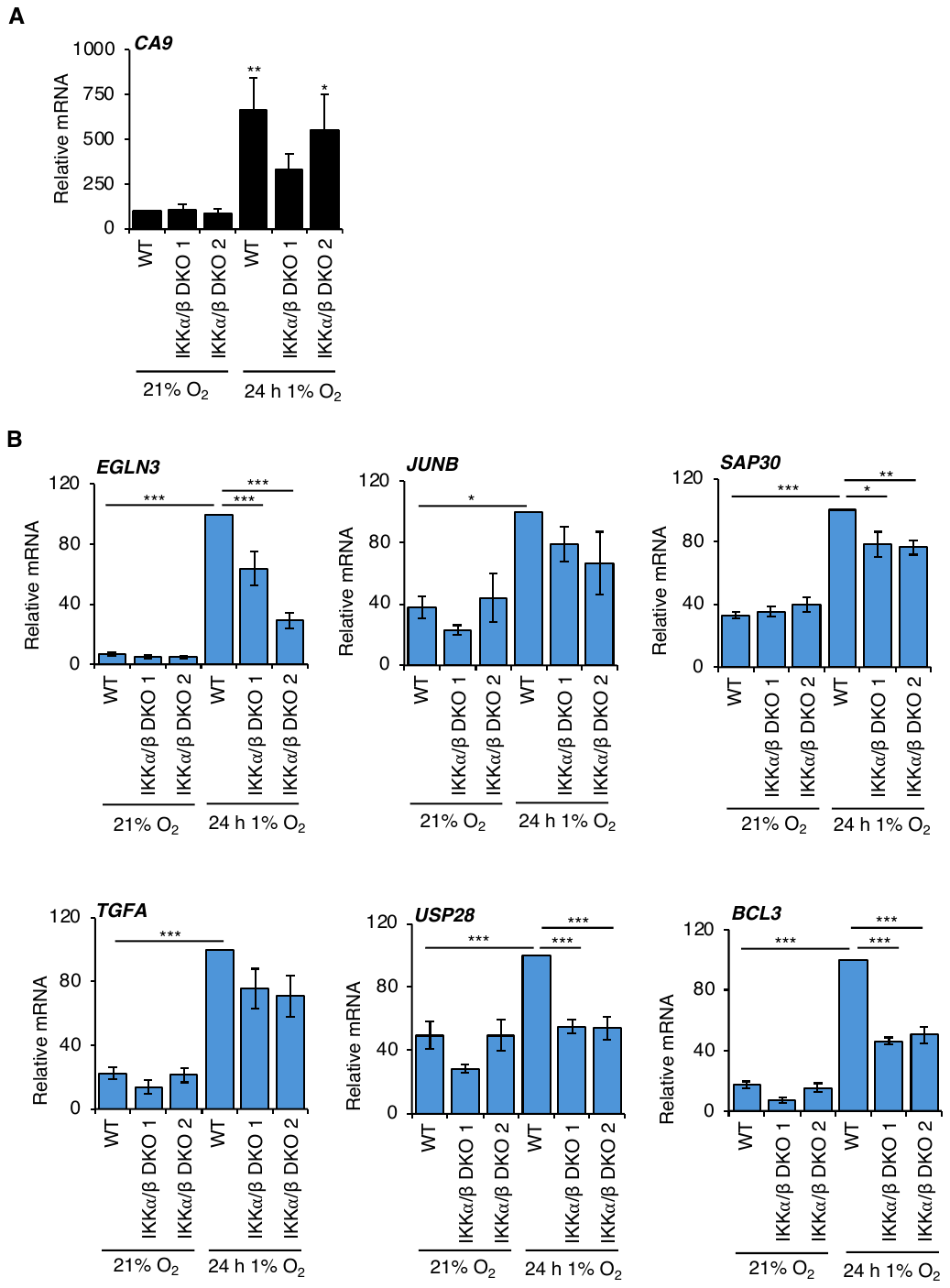
Appendix Figure S17 **NF-𝜅B dependent gene expression changes in HCT116 IKKα/β double knockout CRISPR cell lines in normoxia and hypoxia. A-D.** qPCR analysis in wild type (WT) and IKKα/β double knockout (DKO) HCT116 CRISPR cell lines cultured at 21% oxygen (control) or 24 h 1% oxygen (hypoxia). **D.** *CA9* (**A**), RelA- (**B**), RelB- (**C**), and cRel-dependent (**D**) hypoxia DEGs. Relative mRNA expression levels of the indicated genes were analysed using Actin as a normalising gene. Graphs show mean (n=3) ± SEM, * P < 0.05, ** P < 0.01, *** P < 0.001. **A.** Statistical significance was determined via one-way ANOVA with post-hoc Dunnett’s test (compared to normoxia WT cells). **B-D.** Statistical significance was determined via one-way ANOVA with post-hoc post-hoc Tukey's HSD test.

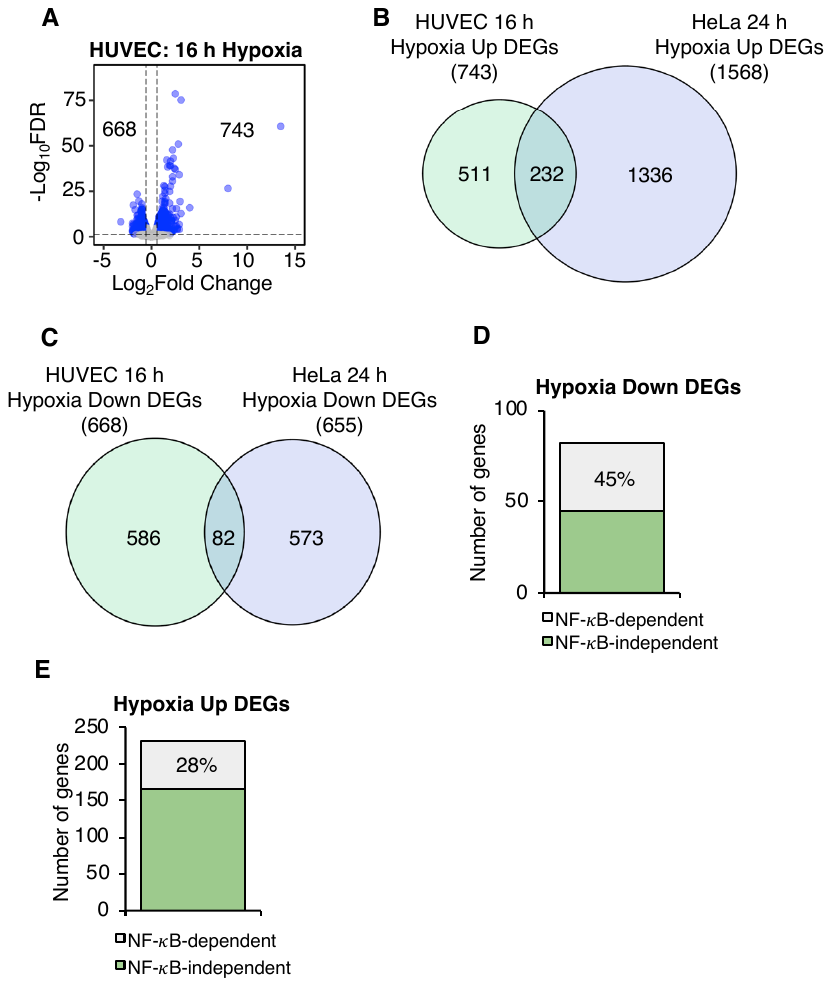

Appendix Figure S18 **Identification of NF-𝜅B dependent hypoxia inducible gene signature in HUVEC cells.**

**A.** Volcano plot displaying differential expression analysis comparing 16 h hypoxia to control from RNA-seq dataset in HUVEC cells. **B-C.** Overlap of hypoxia inducible DEGs identified in HUVEC RNA-seq dataset with HeLa RNA-seq experiment exposed or not to 24 h hypoxia. **D-E.** Percentage of NF-𝜅B dependent hypoxia inducible DEGs are displayed.

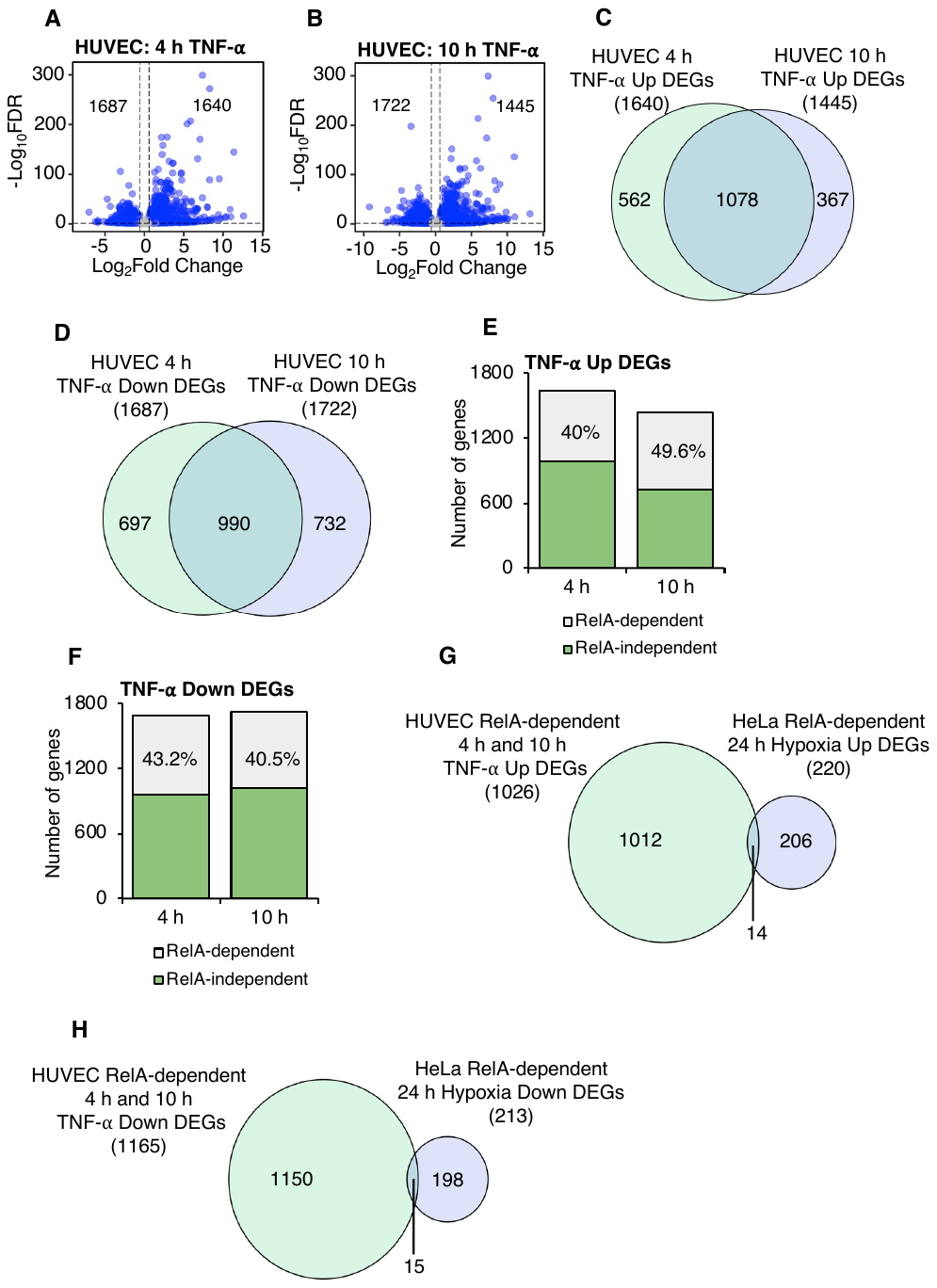
Appendix Figure S19 **Correlation of NF-𝜅B dependent TNF-α inducible gene signature with hypoxia inducible gene signature in HUVEC cells.**

**A-B.** Volcano plots displaying differential expression analysis comparing 4 h (**A**) or 10 h (**B**) TNF-α to control from RNA-seq dataset in HUVEC cells. **C-D.** Overlap of 4 h and 10 h TNF-α inducible DEGs identified in HUVEC RNA-seq dataset. **E-F.** Percentage of RelA-dependent TNF-α inducible DEGs are displayed. **G-H.** Overlap of RelA-dependent TNF-α inducible DEGs in HUVEC cells with hypoxia inducible DEGs identified in HeLa RNA-seq dataset.

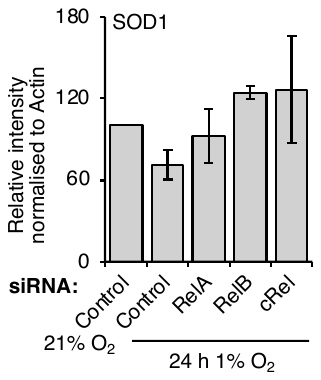

Appendix Figure S20 **Quantification of SOD1 western blot analysis.**

Immunoblot analysis quantification SOD1 in HeLa cells exposed or not to 1% oxygen (hypoxia) for 24 h, with siRNA transfection of control, RelA, RelB or cRel. Signal intensity normalised to Actin (mean n=3, ± SEM) from immunoblot analysis shown in Figure 6C.

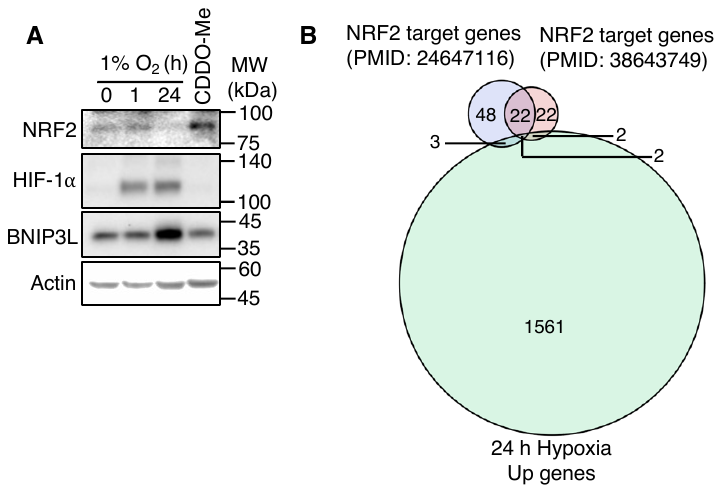

Appendix Figure S21 **Investigation of NRF2 induction in hypoxia.**

**A.** Immunoblot analysis of the indicated proteins in HeLa cells exposed to 1% oxygen for 0,1, and 24 h. Treatment with 100nM of CDDO-Me for 16 h was used as positive control for NRF2 activation. **B.** Overlap of hypoxia upregulated genes with NRF2 target genes.

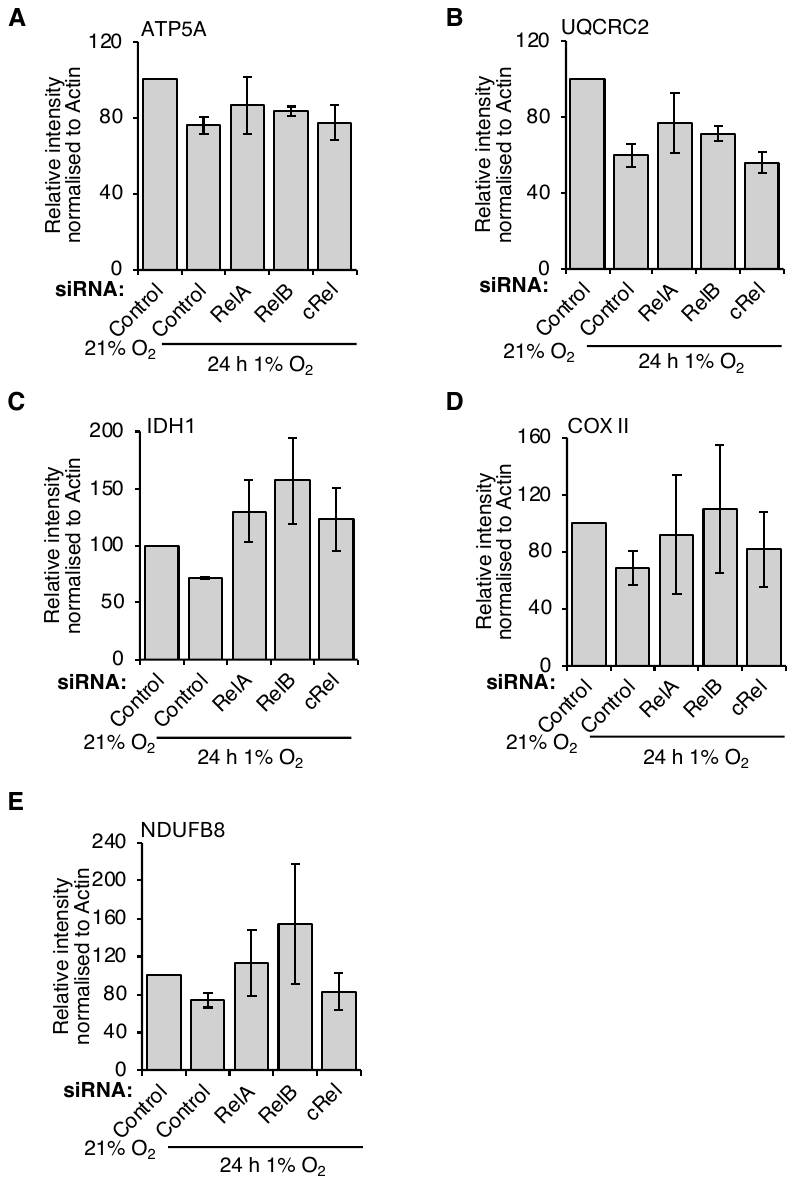

Appendix Figure S22 **NF-𝜅B is required for expression of oxidative phosphorylation related proteins in hypoxia, immunoblot quantifications**

**A-E.** Immunoblot analysis quantification of the indicated proteins in HeLa cells exposed or not to 1% oxygen (hypoxia) for 24 h, with siRNA transfection of control, RelA, RelB or cRel. Signal intensity normalised to Actin (mean n=3, ± SEM) from immunoblot analysis shown in Figure 7.

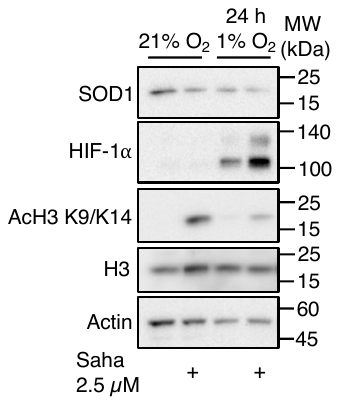

Appendix Figure S23 **SOD1 levels in response to hypoxia and HDAC inhibition.**

Immunoblot analysis of the indicated proteins in HeLa cells exposed or not to 1% oxygen for 24 h, with or without 2.5 µM Saha treatment for 24 h.

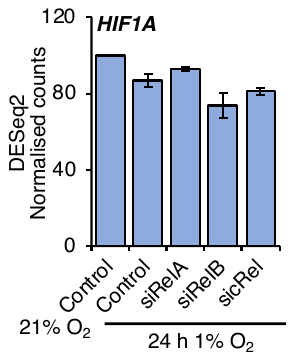

Appendix Figure S24 **HIF-1α RNA levels in response to NF-κB depletion in hypoxia.**

DESeq2 normalised counts acquired from RNA-seq analysis showing differentially expressed HIF1A gene in hypoxia control, and siRelA, siRelB and sicRel with hypoxia, compared to normoxia control.

Appendix Table S1 **NF-𝜅B-dependent hypoxia regulated genes information.**

Selected differentially expressed genes (DEGs) showing NF-𝜅B binding based on overlapping with ChIP Atlas datasets; HIF binding based on overlapping with HIF ChIP following 6 h hypoxia datasets; known NF-𝜅B targets are shown with a tick mark (✔️) based on overlapping with stringent Gilmore laboratory NF-𝜅B targets list; known HIF targets are shown with a tick mark (✔️) based on overlapping with Rocha laboratory HIF targets list.

| **Gene Symbol** | **Gene Name** | **NF-𝜅B binding** | **HIF**  **binding** | **Known**  **NF-𝜅B**  **Target** | **Known**  **HIF**  **target** |
| --- | --- | --- | --- | --- | --- |
| BCL3 | B-Cell Lymphoma 3 | RelA | - | ✔️ | - |
| KMT2E | Lysine Methyltransferase 2E | RelA | HIF-2α, HIF-1β | - | - |
| SAP30 | Sin3A Associated Protein 30 | RelB | HIF-1α, HIF-1β | - | - |
| TGFA | Transforming Growth Factor Alpha | - | - | - | ✔️ |
| EGLN3 | HIF-Prolyl Hydroxylase 3 | - | HIF-1α, HIF-2α, HIF-1β | - | ✔️ |
| USP28 | Ubiquitin Specific Peptidase 28 | RelA, RelB | HIF-1α, HIF-2α, HIF-1β | - | - |
| VIM | Vimentin | RelA, RelB | - | ✔️ | ✔️ |
| NFIX | Nuclear Factor 1X | RelA | HIF-1α, HIF-2α, HIF-1β | - | - |
| GADD45B | Growth Arrest and DNA Damage Inducible Beta | cRel | - | ✔️ | - |
| JUNB | Activator Protein 1/ Jun B Proto-oncogene | cRel | - | ✔️ | - |
| KLF10 | TGFB-Inducible Early Growth Response Protein 1 | cRel | HIF-1β | - | - |
| FTH1 | Ferritin Heavy Chain 1 | RelA, cRel | - | ✔️ | - |
| GCLM | Glutamate-Cysteine Ligase Modifier Subunit | RelA | - | ✔️ | - |
| CCND3 | Cyclin D3 | - | - | - | - |
| NQO1 | NAD(P)H Quinone Dehydrogenase 1 | - | - | ✔️ | - |
| AIFM1 | Apoptosis Inducing Factor Mitochondria Associated 1 | - | - | - | - |
| SOD1 | Superoxide Dismutase 1 | - | - | ✔️ | - |
| CASP10 | Caspase 10, Apoptosis-Related Cysteine Peptidase | RelB, cRel | - | - | - |
| LAMTOR2 | Late Endosomal/Lysosomal Adaptor, MAPK And MTOR Activator 2 | RelB | - | - | - |
| IDH1 | Isocitrate Dehydrogenase (NADP (+)) 1 | RelA | - | - | - |

Appendix Table S2 **Hypoxia RNA-seq dataset information.**

Hypoxia RNA-seq dataset summary

| **Cell line** | **Tissue** | **Control** | **Hypoxia** | **NCBI GEO ID** |
| --- | --- | --- | --- | --- |
| MCF-7 | Breast adenocarcinoma | 21% oxygen | 24 hours,  1% oxygen | GSE153291 |
| A549 | Lung carcinoma |  |  | GSE186370 |
| HCT116 | Colorectal carcinoma |  |  | GSE81513 |
| U87 | Brain glioblastoma |  |  | GSE78025 |
| 501-MEL | Skin melanoma |  |  | GSE132624 |
| SKNAS | Neuroblastoma |  |  | GSE284326 |

Appendix Table S3 **Hypoxia RNA-seq dataset information.**

Sequence Read Archive (SRA) accession numbers for hypoxia RNA-seq datasets.

| **Cell line** | **Control #1** | **Control #2** | **Control #3** | **Hypoxia #1** | **Hypoxia #2** | **Hypoxia #3** |
| --- | --- | --- | --- | --- | --- | --- |
| MCF-7 | SRR12094720 | SRR12094721 | SRR12094722 | SRR12094723 | SRR12094724 | SRR12094725 |
| A549 | SRR16531807 | SRR16531808 | SRR16531809 | SRR16531810 | SRR16531811 | SRR16531812 |
| HCT116 | SRR3535647 | SRR3535651 | SRR3535655 | SRR3535683 | SRR3535687 | SRR3535691 |
| U87 | SRR3175550 | SRR3175551 | SRR3175552 | SRR3175553 | SRR3175554 | SRR3175555 |
| 501-MEL | SRR9283239 | SRR9283240 | SRR9283241 | SRR9283245 | SRR9283246 | SRR9283247 |
